# A novel sustained release therapy of combined VEGF and TNF-α inhibitors leads to pan-ocular protection for months after severe ocular trauma

**DOI:** 10.1101/2023.03.14.531626

**Authors:** Chengxin Zhou, Fengyang Lei, Pui-Chuen Hui, Natalie Wolkow, Claes H. Dohlman, Demetrios G. Vavvas, James Chodosh, Eleftherios I. Paschalis

## Abstract

**Purpose:** To develop a clinically feasible and practical therapy for multi-ocular protection following ocular injury by using a thermosensitive drug delivery system (DDS) for sustained delivery of TNF-α and VEGF inhibitors to the eye.

**Methods:** A thermosensitive, biodegradable hydrogel DDS (PLGA-PEG-PLGA triblock polymer) loaded with 0.7mg of adalimumab and 1.4 mg of aflibercept was injected subconjunctivally in Dutch-belted pigmented rabbits after corneal alkali injury. The polymer was tuned to transition from liquid to gel upon contact with body temperature without need of a catalyst. Control rabbits received 2mg of IgG loaded DDS or 1.4mg aflibercept loaded DDS. Animals were followed for 3 months and assessed for tolerability and prevention of corneal neovascularization (NV), improvement of corneal re-epithelialization, inhibition of retinal ganglion cell (RGC) and optic nerve axon loss, and inhibition of immune cell infiltration into the cornea. Drug release kinetics was assessed *in vivo* using aqueous humor protein analysis.

**Results:** A single subconjunctival administration of dual anti-TNFα/anti-VEGF DDS achieved sustained 3-month delivery of antibodies to the anterior chamber, iris, ciliary body, and retina. Administration after corneal alkali burn suppressed CD45^+^ immune cell infiltration into the cornea, completely inhibited cornea NV for 3 months, accelerated corneal re-epithelialization and wound healing, and prevented RGC and optic nerve axon loss at 3 months. In contrast, anti-VEGF alone or IgG DDS treatment led to persistent corneal epithelial defect, increased infiltration of CD45^+^ immune cells into the cornea, and significant loss of RGCs and optic nerve axons at 3 months. Aqueous humor protein analysis showed first-order release kinetics without adverse effects at the injection site.

**Conclusion:** Sustained concomitant inhibition of TNF-α and VEGF using a biodegradable, slow-release thermosensitive DDS provides significant ocular protection and prevents corneal neovascularization and irreversible damage to retina and optic nerve after corneal alkali injury. This therapeutic approach has the potential to dramatically improve the outcomes of severe ocular injuries in patients.

## Introduction

Ocular injuries can result in irreversible vision loss not only due to corneal damage^[1][2]^ but also due to intraocular complications, such as proliferative vitreoretinopathy (PVR)^[3]^ and secondary glaucoma, the latter a frequent and devastating long-term complication[^4, 5]^. New therapies addressing damage to the anterior and posterior segment from ocular injuries are urgently needed to improve patient outcomes.

To this end, VEGF inhibitors have been proposed as an alternative to corticosteroids for the prevention and treatment of corneal neovascularization (NV)[^6, 7^] in pre-clinical^[8-10]^ and clinical studies^[11-13]^. Likewise, TNF-α inhibitor^[14]^ were shown to provide significant neuroretinal protection against post-injury neuroinflammation in animal studies[^15, 16^]. Moreover, combination therapy with TNF-α and VEGF inhibitors was shown to improve clinical outcomes in patients with age related macula degeneration (AMD) and macular edema[^17, 18^]. Thus, inhibition of angiogenesis and inflammation using antibodies could prove to be an important therapeutic modality for pan-ocular protection after eye trauma.

Antibody delivery to ocular tissues is a challenging task compared to other tissue targets, and may lead to significant adverse events^[19]^. For example, systemic administration of antibodies, especially VEGF inhibitors, only offers limited drug availability in the ocular tissue while exposes the whole body to the agent and can lead to major complications. In an effort to minimize potential adverse effects, topical administration of antibodies in the form of eye drops has been attempted but was not shown to achieve adequate bioavailability in the eye[^20, 21^]. Moreover, one study has shown that prolonged topical therapy with VEGF inhibitors can lead to epithelial toxicity[^12, 22^]. Intravitreal administration of antibodies, on the other hand, achieves excellent bioavailability in the eye but has limited therapeutic effect on the anterior eye and cornea and are associated with rare but devastating complications such as endophthalmitis and retinal detachment. Subconjunctival administration of antibodies is an alternative option allowing good drug bioavailability for both anterior segment and posterior segment[^23, 24^]. However, antibodies undergo rapid diffusion in subconjunctival compartment^[23]^ compared to other competing drug elimination routes, thereby limiting the duration of therapeutic effect.

We addressed these limitations with a thermosensitive biodegradable PLGA-PEG-PLGA triblock hydrogel as a drug delivery system (DDS), and further explore the combined therapeutic efficacy of anti-VEGF and anti-TNF-α given together via a DDS through a single subconjunctival injection. We demonstrate that the thermosensitive DDS can be fine-tuned to transition from liquid to gel at body temperature after injection into the subconjunctival space. This transition process is extremely benign, it involved the displacement of water molecules from the polymer chain, does not require a catalyst, and is not harmful to the antibodies and to the eye. We demonstrate sustained delivery of anti-VEGF and anti-TNF-α to the cornea, iris and retina for 3 months, without local adverse events. Further, using a well-established alkali injury model, we demonstrate almost complete protection from post-injury corneal neovascularization, as well as secondary retinal and optic nerve damage. Our data suggest that sustained delivery of TNF-α and VEGF inhibitors in combination in the appropriate DDS could greatly ameliorate downstream damage from eye injuries.

## Methods

### Thermosensitive polymer solution preparation

To generate a thermosensitive, injectable, biodegradable hydrogel for delivery of therapeutic biologics, a poly(lactide-co-glycolide)-***b***-poly(ethylene glycol)-***b***-poly (lactide-co-glycolide) triblock copolymer was dissolved (15:1 lactic acid: glycolic acid, 1750-1500-1750 Da) (AK141, PolysciTech, West Lafayette, IN, USA) in sterile water (20% w/v) at 4°C overnight with gentle stirring. The polymer solution was then sterilized with 30 minutes ultraviolet light radiation in a biosafety hood. The sterile polymer solution was stored in syringes in -20°C until use.

To prepare the drug-loaded polymer solution, infliximab lyophilized powder (Janssen biotech, Raritan, New Jersey, USA) was weighed and dissolved in the sterile polymer solution at 4 °C. The aflibercept injection solution (Regeneron, Tarrytown, NY, USA) was added to the solution by at 4 °C. The final product contained 2mg of therapeutic proteins (1:2 infliximab:aflibercept) in every 300μL of polymer solution (refilled in a 1mL syringe).

### Rheological characterization

Rheological properties of the 20% triblock polymer were measured by dynamic mechanical analyzer (Discovery HR-3, TA instruments) with a 20mm, 5 degree cone. The test was performed by oscillating at an angular frequency 6.283 rad/s, 0.1% strain, in increments of 1°C ranging from 4-45°C with 1 minute of temperature equilibration at each temperature.

### Corneal neovascularization model

A corneal alkali injury was used to generate corneal NV, as previously published[^14-16, 25-29^]. All rabbits were treated in accordance with the Association for Research in Vision and Ophthalmology Statement on the Use of Animals in Ophthalmic and Vision Research and the National Institutes of Health Guide for the Care and Use of Laboratory Animals, and the experimental protocol was approved by the Animal Care Committee of the Massachusetts Eye and Ear. Ten female Dutch-Belted rabbits (2 - 2.5kg), (Envigo, Dedham, MA, USA) were anesthetized by intramuscular injection of ketamine hydrochloride INJ, USP (35mg/kg; KetaVed, VEDCO, St. Joseph, MO, USA) and xylazine (5mg/kg; AnaSed, LLOYD, Shenandoah, IA. USA). Topical anesthetic (0.5% proparacaine hydrochloride, Bausch & Lomb, Tampa, FL, USA) was applied to the eye. An alkali injury was created using an 8-mm diameter cotton sponge soaked in 2N NaOH, applied to the cornea for 20 seconds. Following removal of the filter paper, the eye was immediately irrigated with normal saline solution for 15 minutes.

### Subconjunctival DDS placement

Each 20% triblock polymer was UV-radiated for 30 mins on ice, and mixed with either a pre-determined amount of Infliximab dry powder, aflibercept stock solution, or both. The drug-laden triblock polymer solution was distributed in 1-mL syringes for in vivo injection. Each syringe was pre-filled with 300μL of drug-laden DDS and was stored at 4 °C until animal injection in vivo.

For DDS administration, chilled DDS (300μL/syringe/injection) loaded either with 2mg of infliximab/aflibercept antibodies (1:2), 1.3mg of aflibercept (*n* = 3), or 2 mg of human IgG isotype (*n* = 4) (I4506-100MG, Millipore-Sigma, Saint Louis, MO,USA) were injected subconjunctivally in the upper eyelid of eyes immediately after the post-alkali exposure irrigation, i.e., 15 minutes after the injury. Erythromycin ophthalmic ointment (0.5%, Bausch & Lomb) was administered topically twice a day for 1 week.

### Evaluation of corneal NV and epithelial defects

Evaluation of corneal NV was performed under general anesthesia every week for the first month and every two weeks thereafter for two additional months. All treated and control eyes were photographed using a digital SLR camera (Nikon, Tokyo, Japan) attached to a surgical microscope (S21; Carl Zeiss, Jena, Germany) at standard magnifications. Corneal epithelial defects were stained with fluorescein and imaged using a portable slit-lamp (Keeler 3010-P-2001; Keeler Americas, Malvern, PA, USA) equipped with cobalt blue filter and a mounted digital camera at magnification ×10. Photographs were analyzed using ImageJ 1.50e software (http://imagej.nih.gov/ij/; provided in the public domain by the National Institutes of Health [NIH], Bethesda, MD, USA). For any epithelial defect, the total corneal area stained with fluorescein (pixel2) was normalized relative to the whole cornea area (pixel2) in the same image, yielding the corneal epithelial defect area per whole cornea ratio (%). For analysis of corneal NV, each cornea was divided into superior and inferior halves. The vascularized area in each half was quantified and normalized to half of the corneal area (%). Illustration graphs of all the representative biomicroscopic images of corneal neovascularization and epithelial defects were manually sketched in Adobe Illustrator 2020 software (Adobe, San Jose, CA) by delineating the corneal vessels and defects with a ‘pencil’ tool on top of the original photos.

### ELISA assay

Rabbits aqueous humor (AH) was collected when animal was under general anesthesia at baseline, weekly for 1 month, and bi-weekly for additional 2 months post injection. Topical anesthetic (0.5% proparacaine hydrochloride, Bausch & Lomb, Tampa, FL, USA) was applied to the eye. A 30-G needle combined with a 1-mL syringe was used to aspirate approximately 50∼70μL aqueous humor from the treated eye and the contralateral eye. Antibiotics eye drops were given to the eye following the procedure. Aqueous humor samples were stored in -80 °C freezer until processed with ELISA assay. All AH samples were diluted in PBS at 1:20 and stored on ice. Human IgG ELISA assay (RAB0001, Sigma, Saint Louis, MO) was performed per manufacturer’s protocol. Serially diluted human IgG standards and blank samples were analyzed along with the test samples. A standard curve was generated to calculate the human IgG concentrations in the AH.

### Histological and immunohistochemical evaluation of the cornea

At three months after injury, rabbits were euthanized using intravenous Beuthanasia-D (sodium pentobarbital and phenytoin sodium, 100 mg/kg; Merck Animal Health, Madison, NJ, USA). Tissues were fixed in 4% paraformaldehyde (PFA and embedded in optimal compound temperature (OCT). Tissue frozen sections (10 µm thickness) were prepared using a cryostat (CM1950; Leica Biosystems, Buffalo Grove, IL, USA). Hematoxylin and eosin (H&E) staining was performed on methacrylate-embedded tissue sections for histologic evaluation. Immunohistochemical evaluation was performed as previously described.^[30]^ Primary antibodies diluted in 1% BSA were incubated with the tissue samples overnight at 4°C, followed by secondary antibody incubation for two hours at room temperature. Detail information of the antibodies is provided in the Supplementary Methods section. Immune cell recruitment and cytokine expression were assessed using antibodies against CD45 (1:100, SC-70690, Santa Cruz, Dallas, TX, USA). Residual human IgG in the DDS and in the ocular tissues was determined using goat anti-human IgG antibody (A-21089, Thermo Scientific, Waltham, MA, USA). The total number and area of cells expressing CD45 in corneal sections were determined using the “Analyze Particles” tool in ImageJ software, as previously described.^[19]^ Mean ±standard deviation (SD) of different rabbits were reported.

## Results

### DDS s*ol-gel* transition at body temperature

The PLGA-PEG-PLGA triblock copolymer used in this study was tuned to exhibit peak storage and loss moduli at a sol-gel transition temperature of ∼37°C (body temperature) (**Fig.1 A-B, D**). This process is achieved by a thermosensitive reaction that displaces water molecules from the triblock PEG-PLGA-PEG polymer chain, which in turn causes the liquid polymer to transition to gel without the need to a catalyst. The process is totally benign and does not cause damage to the antibodies or the tissue. Cured 20% triblock hydrogel remains viscus and transparent after gelation.

**Figure 1.**
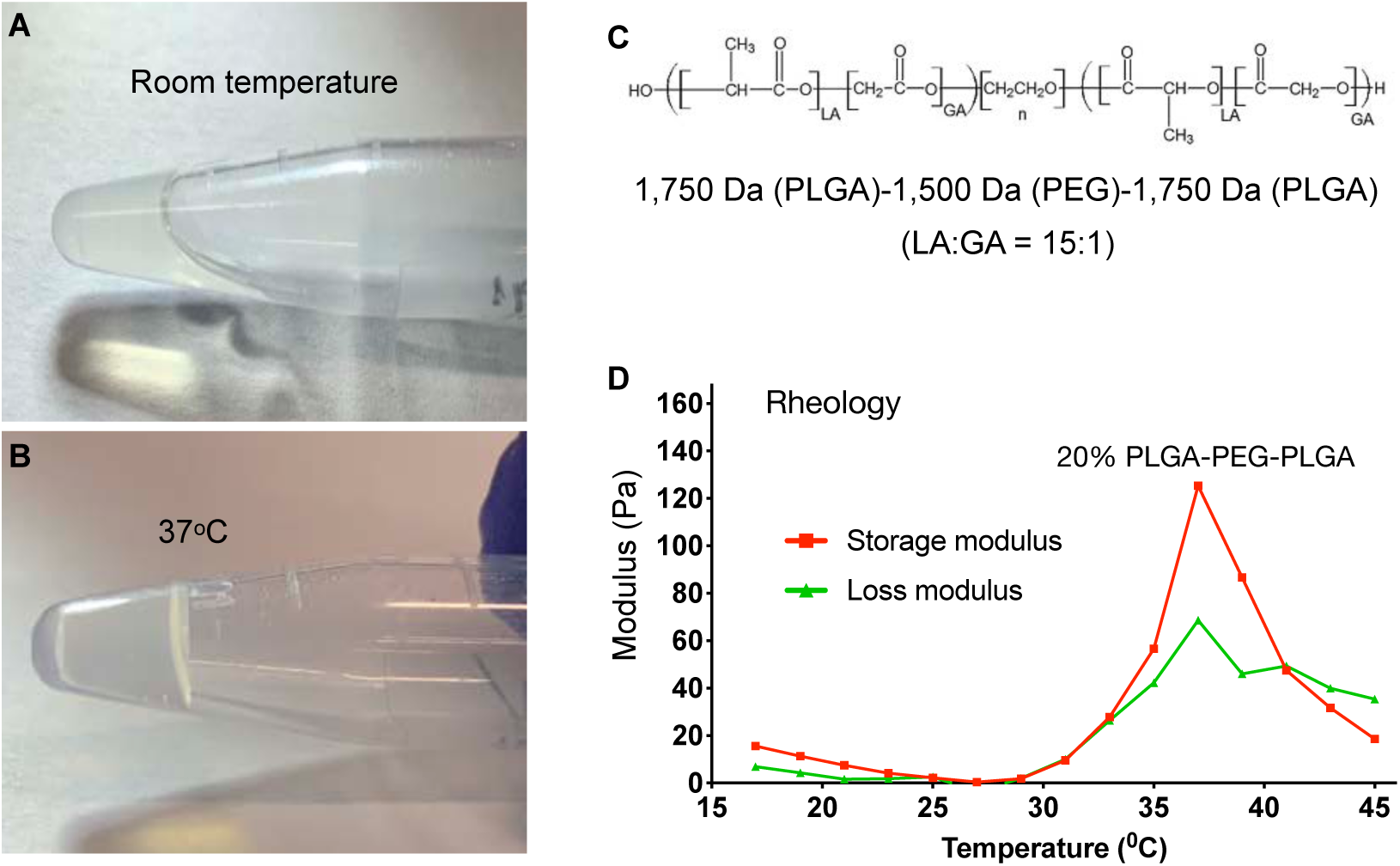
Characterization of 20% PLGA-PEG-PLGA triblock polymer. (**A-B**) Sol-gel transition from room temperature to 37°C. Note the liquid polymer becomes a gel. (**C**) Molecular structure of the triblock polymer and (**D**) rheological assessment of the triblock polymer showing sol-gel transition peak at 37°C.

### Long-term sustained release of antibodies into the eye after single subconjunctival injection of the DDS

The ability to deliver anti-VEGF/anti-TNF-α inhibitors, as well as the safety and efficacy of the thermosensitive triblock polymer was evaluated *in vivo*, following a single subconjunctival injection of the DDS in rabbit eyes with corneal alkali injuries. The DDS was injected as cold liquid polymer in the superior bulbar conjunctival 15 minutes after the corneal alkali burn using a 30G needle and a 1-mL syringe. Upon injection, the polymer rapidly gelated (**Fig. 2 A**) and formed a visible drug reservoir. The DDS biodegraded over the 3 months of follow-up (**Fig. 2 B-G**). Antibody penetration in the eye was evaluated by aqueous humor sampling and analysis of IgG content using a human IgG ELISA kit (RAB0001, Sigma, Saint Louis, MO). Upon DDS injection, human IgG content in the aqueous human exhibited a rapid increase for the first two weeks after injection (**Fig. 2 H**), followed by gradual decrease between weeks 2-6 and subsequent normalization between weeks 6-12 (**Fig. 2 H**).

**Figure 2.**
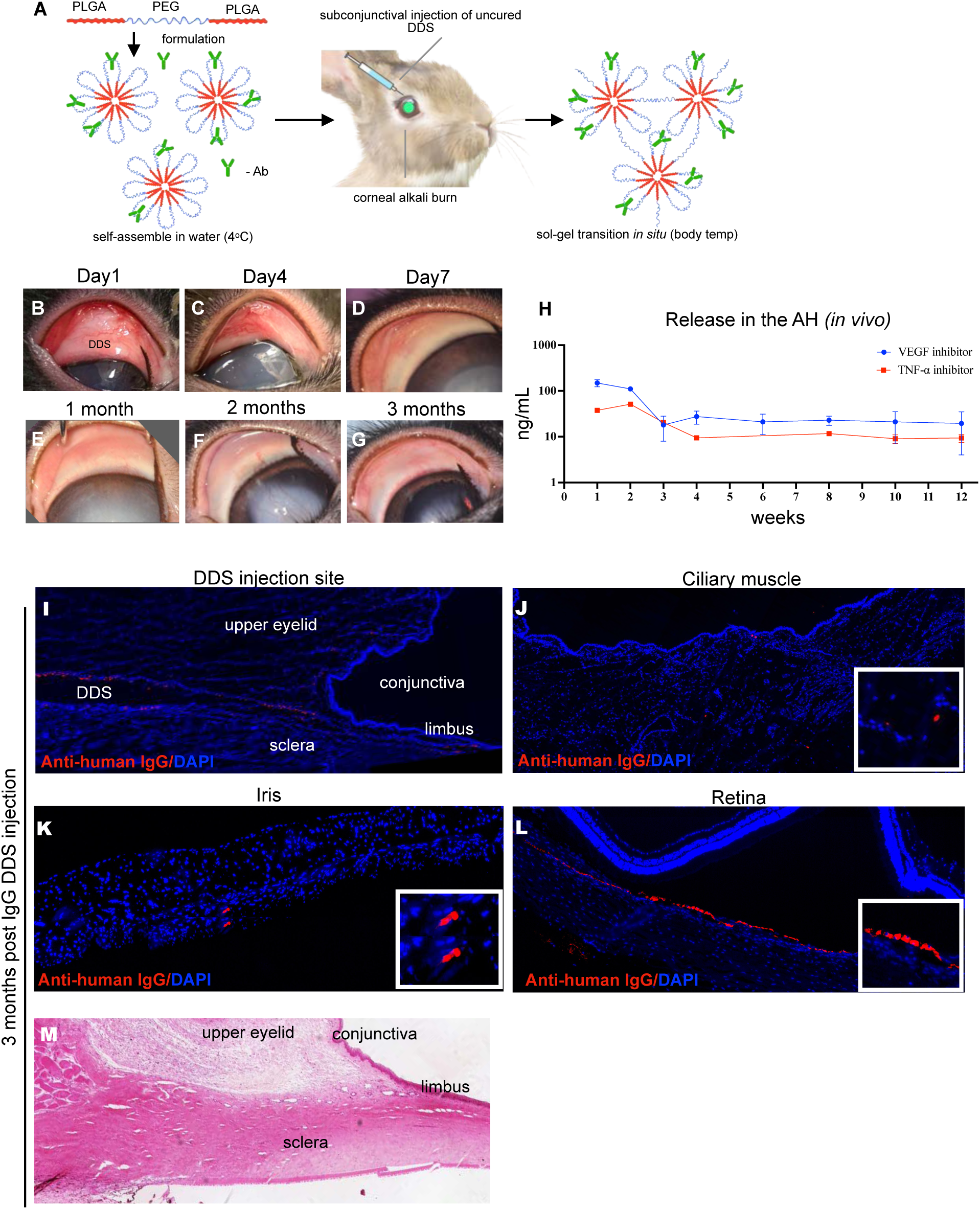
*In vivo* evaluation of drug release following subconjunctival administration of the 20% triblock thermosensitive drug delivery system (DDS). (**A**) Schematic representation of the DDS containing antibodies injected in the superior subconjunctival space after corneal alkali burn. (**B-G**) Biomicroscopic images of the upper conjunctiva and cornea harboring the DDS. The DDS slowly degraded, as evident by the regression of upper conjunctival thickening over 3 months. (**H**) ELISA analysis of aqueous humor samples demonstrated continuous release of therapeutic levels of antibodies (anti-TNF-α and anti-VEGF) in the anterior chamber for over 3 months. Three months after DDS injection, (**I**) the subconjunctival tissue harboring the DDS is immunopositive for human IgG, suggesting presence of humanized antibody in the tissue. Likewise, humanized antibody is also found in the ciliary muscle **(J)**, iris **(K)**, and (sub-retinal space **(L)**. (**M**) H&E staining of upper eyelid hosting the DDS showed no tissue abnormalities. The DDS appeared to be degraded and cleared from the tissue at 3 months.

Immunofluorescent staining of human IgG in cryosectioned eyelid tissues showed that 3 months after injection of the DDS, a substantial amount of human IgG was present in the bulbar and forniceal subconjunctival connective tissue, iris, ciliary body, and retina (**Fig. 2 I-L**). Histopathological examination 3 months after injection revealed normal-appearing ocular adnexa (**Fig. 2 M**).

### Anti-TNFα/anti-VEGF DDS treatment completely suppresses corneal angiogenesis after injury

Central corneal alkali burn in rabbit eyes led to progressive corneal NV in the superior and inferior corneal regions of IgG DDS treated eyes **(Fig. 3 A**). Corneal NV reached its peak at 1.5 months post injury (25% of superior cornea; 30% of inferior cornea) **(Fig. 3 A-D**). In comparison, anti-VEGF DDS treatment led to partial suppression of NV at 3 months, especially in the inferior cornea (maximal inferior NV : **30%** with IgG *vs*. **8%** with anti-VEGF. \**P<0*.*05; Mixed ANOVA test with Tukey’s correction*), but did not completely halt progression, as the area of NV continued to grow in all rabbits during the 3 month follow-up **(Fig. 3 A-D**). Moreover, anti-VEGF DDS treatment showed marginal inhibition of superior corneal NV, as compared to IgG control group (*P >0*.*05; Mixed ANOVA test with Tukey’s correction)*. In contrast, anti-TNFα/anti-VEGF DDS treatment led to complete inhibition of NV in the superior and inferior cornea for 3 months, as compared to the IgG DDS treatment group (\**P*<0.05; \*\*\**P*<0.001; Mixed ANOVA test with Tukey’s correction), **(Fig. 3 A-D).**

**Figure 3.**
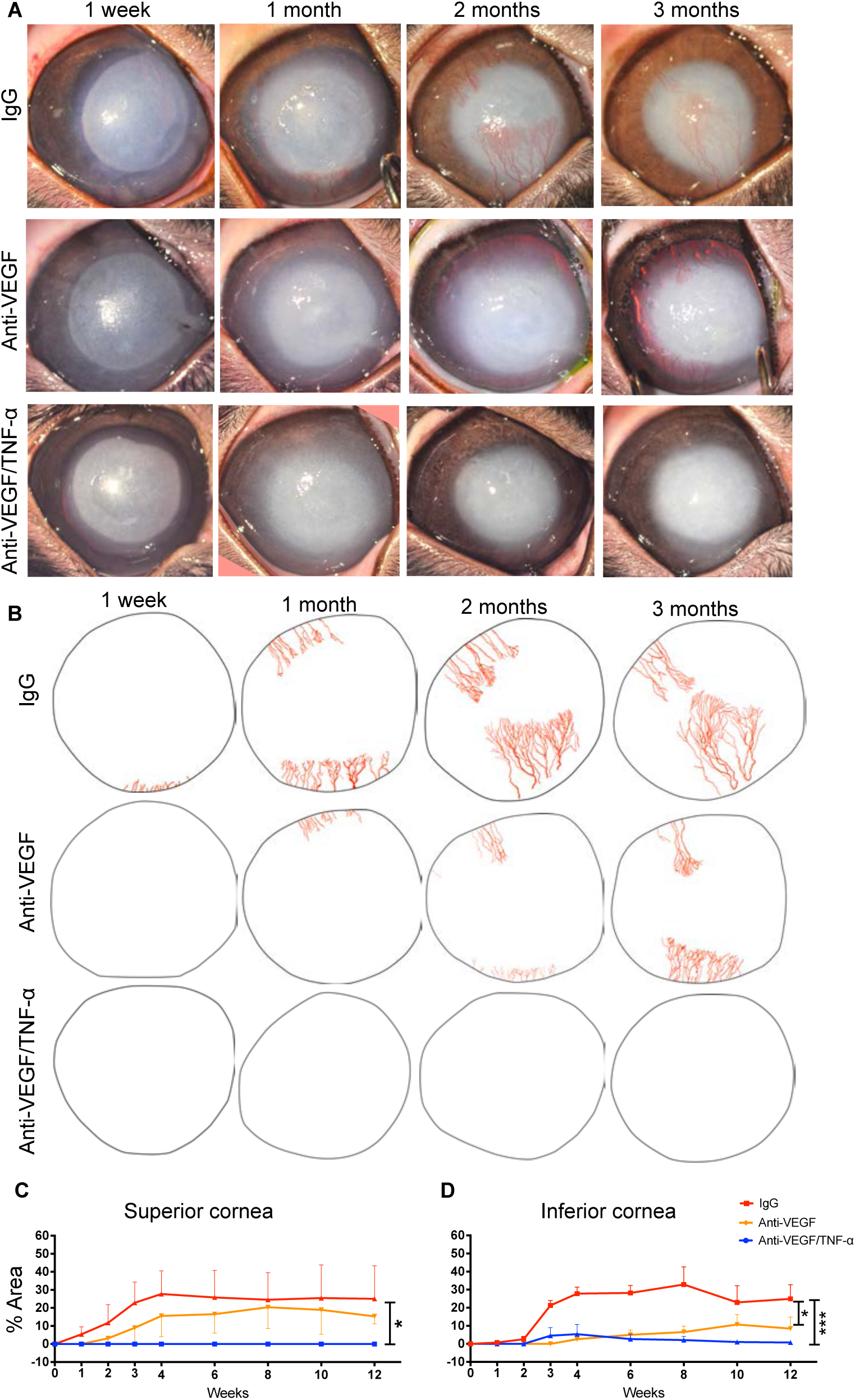
Anti-TNFα/anti-VEGF DDS antibodies sustained released by thermogel DDS significantly halted the progression of corneal neovascularization after corneal burn injury. (**A**) Representative biomicroscopic images of burned, DDS-implanted rabbit eyes at specified time points. (**B**) Angiographic illustration of the corneas presented in (**A**). (**C-D**) CoNV area quantification in percentages of superior and inferior cornea areas, respectively. (**A-D**) Three months after injury, anti-TNFα/anti-VEGF DDS conferred nearly complete blockade of CoNV (**C, D**; *Blue curve*). In contrast, eyes administrated with isotype IgG DDS exhibited extensive CoNV over 3 months (CoNV area at endpoint: ∼30% of superior area; 30% of inferior cornea) (**C**,**D**; *Red curves*). Eyes treated with aflibercept DDS showed milder yet progressive CoNV when compared to the IgG DDS eyes during the 3-months (CoNV area at endpoint:∼15% of superior cornea; ∼8% of inferior cornea) (**C**,**D**; *Orange curves*). Mixed ANOVA. *\*P<0*.*05; ***P<0*.*001*.

### Anti-TNFα/anti-VEGF DDS treatment improves corneal epithelial healing after injury

IgG DDS treated eyes had persistent epithelial defects even at 1 month after injury (10% of cornea area), which gradually reduced in subsequent months (**Fig. 4 A-C**). In eyes treated with anti-VEGF DDS, persistent epithelial defects peaked at 2 months after injury (15% of cornea area) without resolution through 3 months (**Fig. 4 A-C**). In contrast, combined anti-TNFα/anti-VEGF DDS treatment led to reduced corneal epithelial defects at 1 month and complete corneal re-epithelialization at 2 months, with a stably intact epithelium when measured at 3 months **(Fig. 4 A-C)**. Quantification of the area of epithelial defect confirmed the effect of combination anti-TNFα/anti-VEGF DDS as compared to the other 2 treatments, (**Fig. 4 C**) (\**P<0*.*05*, mixed ANOVA test with Tukey’s multiple comparison test).

**Figure 4.**
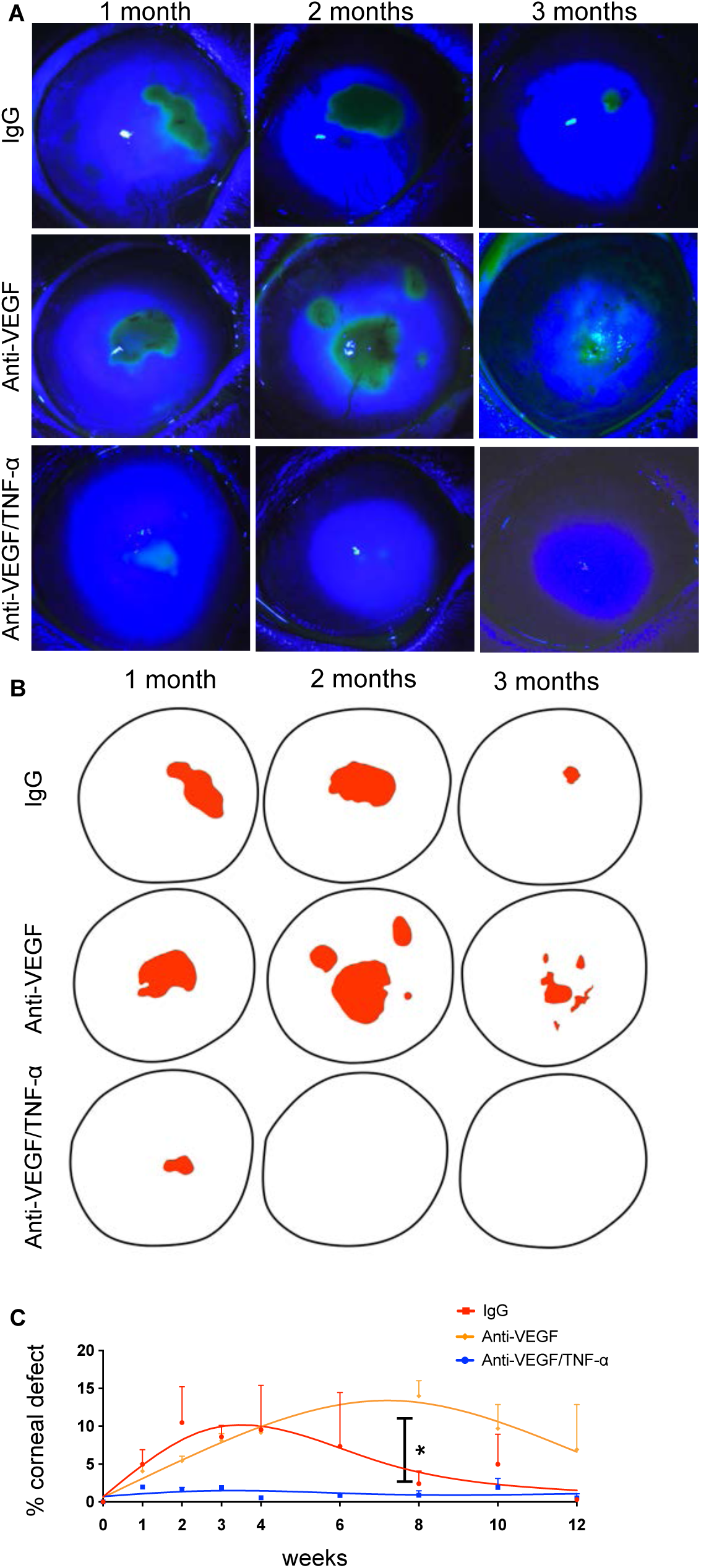
Anti-TNFα/anti-VEGF DDS treatment reduced corneal epithelial defect in the injured cornea. (**A**) slit lamp biomicroscopy of fluorescein stained rabbit eyes implanted with different DDS. (**B**) Image reconstruction of corneal epithelial defects in the burned, DDS-implanted corneas shown in (**A**). (**C**) Corneal epithelial defect quantification in percentage of total corneal area. Anti-TNFα/anti-VEGF DDS treatment significantly suppressed the development of epithelial defect on the burned cornea, as compared to the Aflibercept group and IgG groups. \**P<0*.*05*. Mixed ANOVA test with Tukey’s multiple comparison test.

As shown by immunofluorescent staining of rabbit corneas, combined anti-TNFα/anti-VEGF DDS treatment significantly reduced CD45^+^ immune cell infiltration into the injured cornea (**Fig. 5 C, D**), as compared to IgG DDS (**Fig. 5 A, D**) or anti-VEGF DDS treatment (**Fig. 5 B, D**) (\**P<0*.*05*, one-way ANOVA test with Tukey’s multiple comparison test).

**Figure 5.**
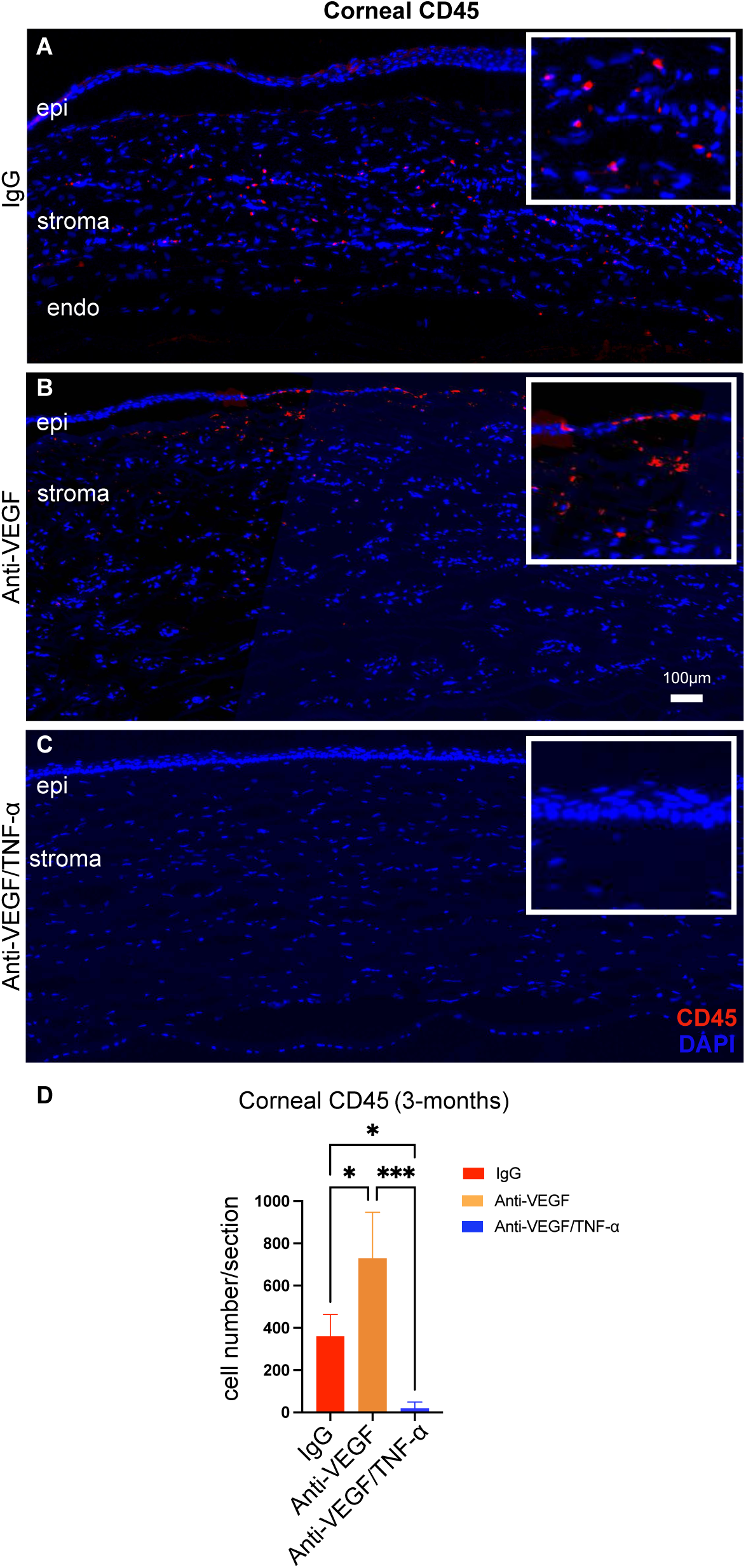
Corneal inflammation and leukocyte infiltration. (**A**) Marked CD45+ cell accumulation in the cornea of IgG (**A**) or Aflibercept (**B**) DDS treated eyes at 3 months of injury. (**C and G**) Remarkable reduction in CD45+ cell accumulation in the cornea following subconjunctival injection of anti-TNFα/anti-VEGF DDS. (**D, E, F)** Immunofluorescent staining of rabbit corneas exhibited limited immunopositivity of TNFα in the central anterior stroma, adjacent to the burn area at 3 months of injury, suggesting the cornea may have transitioned from acute inflammation to tissue regeneration at 3 months. (**H**) Quantification of the TNFα staining area did show statistical significance between the three experimental groups. One-way ANOVA with Tukey’s correction. \**P<0*.*05*.

### Anti-TNF-α/anti-VEGF DDS treatment prevents retinal and optic nerve damage after injury

We previously demonstrated that corneal alkali injury is associated with secondary retinal and optic nerve degeneration in mice, rabbits, and humans[25], and at least in experimental models, that this damage is mediated by secondary inflammation independently of the intraocular pressure[25]. In alkali-injured eyes, combined anti-TNF-α/anti-VEGF DDS treatment significantly reduced post-injury retinal ganglion cell loss at 3 months, as compared to the IgG DDS treatment (2.7% RGC loss vs. 45% RGC loss, respectively; *P*<0.01; **Fig. 6 A-D**). RGC loss was similar between the anti-VEGF and IgG DDS treated groups (63% RGC loss vs. 45% RGC loss, respectively; *P*>0.05. **Fig. 6 B-D**). Combined anti-TNF-α/anti-VEGF DDS treatment also significantly reduced peripheral and central optic nerve axon loss, as compared to IgG (**Fig. 6 E,H, G,I, K**) and anti-VEGF DDS treatments **(Fig. 6 E-L**) (\**P*<0.05; \*\**P*<0.01; \*\*\**P*<0.001; \*\*\*\**P*<0.0001, mixed ANOVA test with Tukey’s correction).

**Figure 6.**
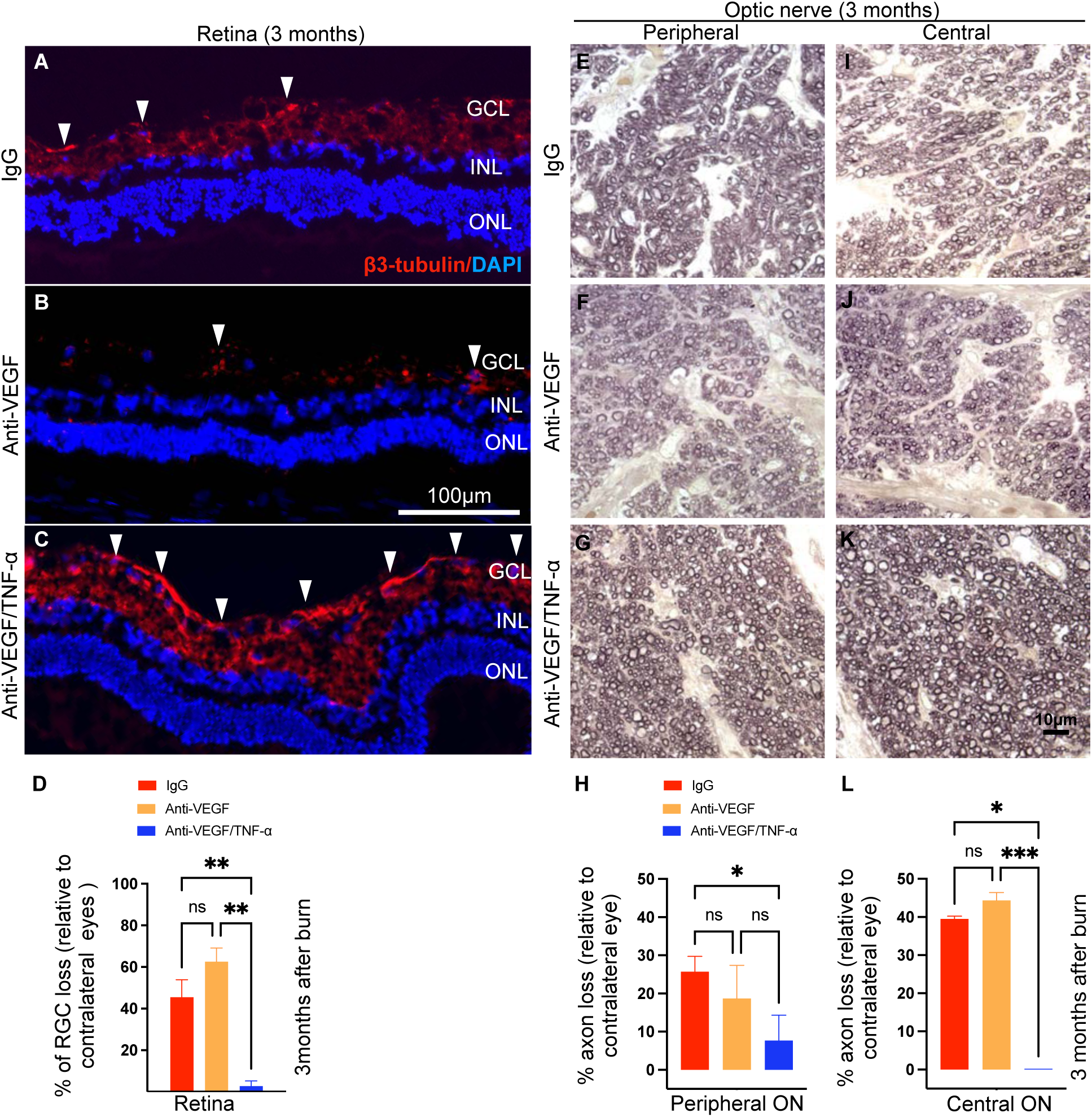
Anti-TNF-α/anti-VEGF DDS treatment effectively ameliorated retinal neuropathy and optic nerve degeneration in the injured eyes. (**A-C, D**) Three months after corneal burn, IgG or aflibercept DDS treated rabbits exhibit significant retinal ganglion cell loss, as indicated by a retinal ganglion cell marker β3-tubulin (red color). In contrast, the anti-TNFα/anti-VEGF DDS provided almost complete protection against RGC loss. ***Arrow head***: a normal ganglion cell. (**E-G**) Representative PPD staining of peripheral rabbit optic nerves (*63X obj*). (**H**) Loss of normal nerve axon loss in peripheral optic nerves relative to the contralateral intact optic nerve. (**I-K**) Representative PPD staining of central rabbit optic nerves (*63X obj*). (**L**) Loss of normal nerve axon loss in central optic nerves relative to the contralateral intact optic nerve. (**E-L**) The protective effects of the anti-TNFα/anti-VEGF DDS was confirmed with PPD staining of the optic nerves, which otherwise showed marked axonal degeneration in the IgG and aflibercept DDS treated eyes and almost complete retention of the nerve axons in anti-TNFα/anti-VEGF DDS treated eyes. One-way ANOVA with Tukey’s correction. *\*P<0*.*005. **P<0*.*01. ***P<0*.*001. ****P<0*.*0001. GCL=ganglion cell layer. INL=inner nuclear layer. ONL=outer nuclear layer. RGC= retinal ganglion cell*.

## Discussion

In the experiments shown herein, we leveraged a tunable, thermosensitive PLGA-PEG-PLGA triblock co-polymer to achieve sustain delivery of VEGF and TNF-α inhibitors to the cornea, iris, and retina for 3 months. Combined treatment with antibodies to both VEGF and TNF-α completely inhibited corneal NV and remarkably reduced retinal and optic nerve damage after corneal alkali injury. Moreover, combined treatment limited a known critical adverse effect of anti-VEGF therapy, specifically the retardation of corneal epithelialization and wound healing[^12, 19^], as manifested by persistent epithelial defect^[31]^.

Previous studies in our group have focused on developing a non-biodegradable macro-porous DDS for anti-TNF-α or anti-VEGF therapy to the eye[^26, 29, 32^]. These studies required placement of the DDS in the subconjunctival space using an incision in the conjunctiva. Here we advance this technology and leverage on a biodegradable thermosensitive PLGA-PEG-PLGA triblock co-polymer able to administer VEGF and TNF-α inhibitors to the eye through a single subconjunctival injection with a 30G needle. The DDS is delivered in liquid phase and becomes a gel upon contact with the tissue at body temperature. Gelation requires no catalysts and does not generate toxic byproducts. Our studies, performed in live animals, confirmed a first-order release kinetics for 3 months in the aqueous humor of rabbits and delivery of therapeutic levels of antibodies to the cornea, iris, and retina for 3 months. Notably, this resulted in not only complete inhibition of corneal NV but reduced retina and optic nerve damage after alkali injury to the eye, a smoldering injury notoriously difficult to mitigate with current ophthalmic treatments[^4, 5, 14, 25, 33, 34]^.

In contrast to single-target therapy with either VEGF or TNF-α inhibitors, concomitant inhibition of TNF-α and VEGF, as demonstrated in this study, is therapeutically superior in preventing cornea neovascularization after alkali injury and minimizing the risk of ocular complications from anti-VEGF administration to the eye[^31, 35, 36^]. Such therapeutic outcome may not be entirely attributed to the anti-angiogenic effect of VEGF inhibitor, but rather to a synergic effect of the TNF-α inhibitor. This is supported by current data showing that aflibercept DDS (treatment control) did not achieve the same level of anti-neovascularization of the cornea and in fact led to persistent epithelial defect that was not present in the combined regimen. Likewise, neither anti-VEGF DDS nor anti-TNF-α DDS therapy alone, as demonstrated in published literature, were able to completely inhibit long-term corneal NV in rabbit corneal alkali injury[^19, 29^]. Collectively, data suggest that simultaneous inhibition of TNF-α and VEGF is more effective than either therapy alone[^26, 37-42^]. Noteworthy, concomitant inhibition of VEGF and TNF-α was previously shown to improve clinical outcomes in AMD and macular edema patients[^17, 18^] confirming our observations in this study and suggesting that concomitant inhibition of VEGF and TNF-α could be a preferred therapy for ocular burns.

Importantly, the proposed therapy offers also significant protection to the neuroretina and optic nerve. A single administration of the anti-TNF-α/anti-VEGF DDS led to dramatical inhibition of retinal ganglion cells and optic nerve axon loss at 3 months after ocular alkali burn, an injury known to result in severe retinal pathology in humans and animals[^5, 25, 34, 43, 44]^. In previous studies we showed that TNF-α is the primary mediator of retinal and optic nerve damage[^16, 25, 26, 28, 29^], through the modulation of neuroglia phenotype[^15, 27^]. Here we show that administration of a total dose of 0.7mg of TNF-α inhibitor results in almost complete retinal and optic nerve protection for 3 months, which is an important improvement to previous studies that used high dose systemic TNF-α inhibitors. We expect that this treatment modality can substantially reduce the risk of systemic exposure to the drug without compromising the therapeutic outcomes to the eye.

In summary, this is the first study to show multi-ocular protection after alkali injury using a single administration of a DDS that delivers TNF-α and VEGF inhibitors to the cornea, iris, and retina for 3 months. The proposed combination therapy is therapeutically superior to anti-VEGF therapy alone against post-injury corneal NV and offers improved retinal and optic nerve protection with minimum risk of systemic exposure to the antibodies. These results may have practical clinical implications for the treatment of patients with ocular injuries, and further clinical studies are warranted.

